## Supporting figures for "Analysis of the *Leishmania mexicana* promastigote cell cycle using imaging flow cytometry provides new insights into cell cycle flexibility and events of short duration"

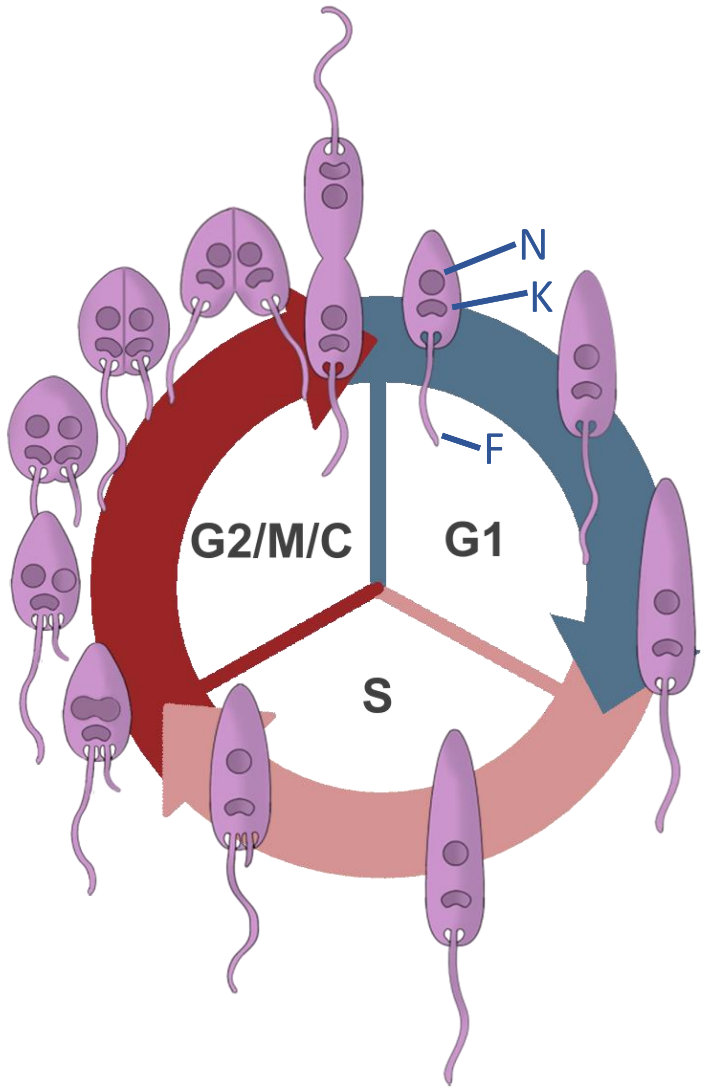


**S1 Fig**

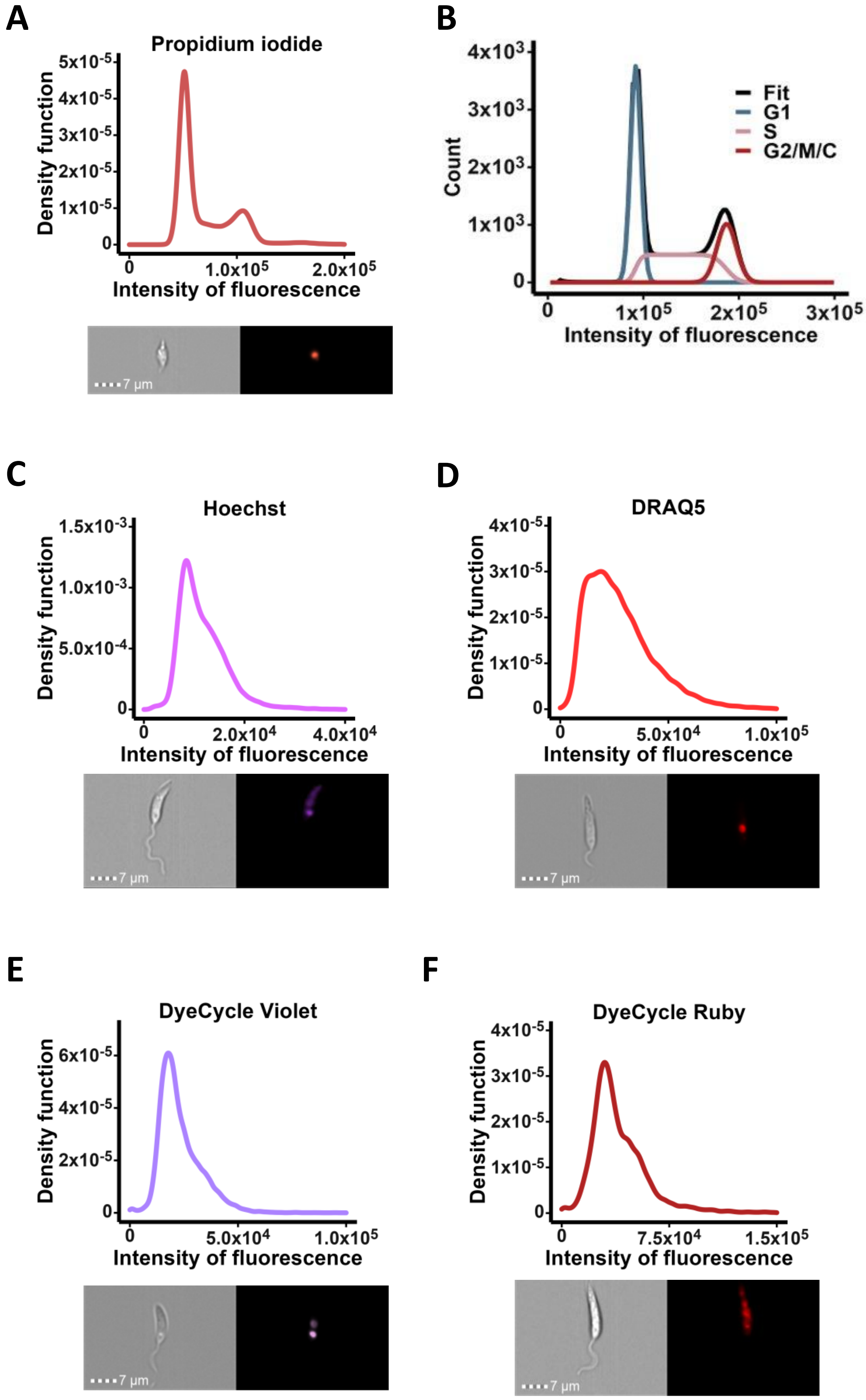
A
Propidium iodide
Density function
Intensity of fluorescence
7 µm
B
Count
Fit
G1
S
G2/M/C
Intensity of fluorescence
C
Hoechst
Density function
Intensity of fluorescence
7 µm
D
DRAQ5
Density function
Intensity of fluorescence
7 µm
E
DyeCycle Violet
Density function
Intensity of fluorescence
7 µm
F
DyeCycle Ruby
Density function
Intensity of fluorescence
7 µm

### S2 Fig

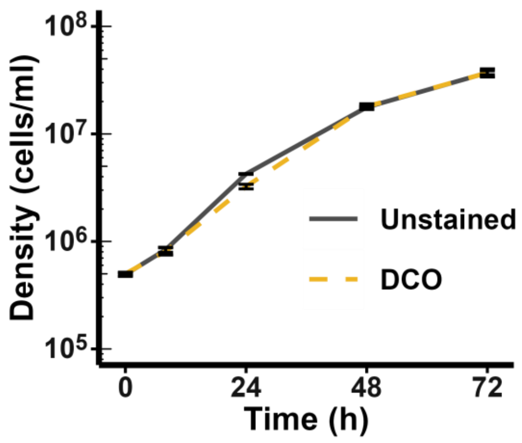

$10^8$
$10^7$
$10^6$
$10^5$
Density (cells/ml)
0
24
48
72
Time (h)
Unstained
DCO


S3 Fig

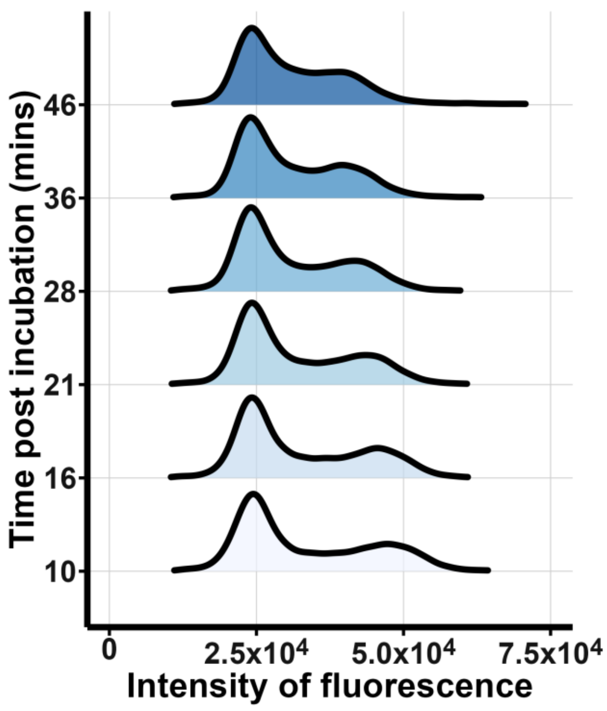
Time post incubation (mins)
46
36
28
21
16
10
0
2.5x10^4
5.0x10^4
7.5x10^4
Intensity of fluorescence

**S4 Fig**

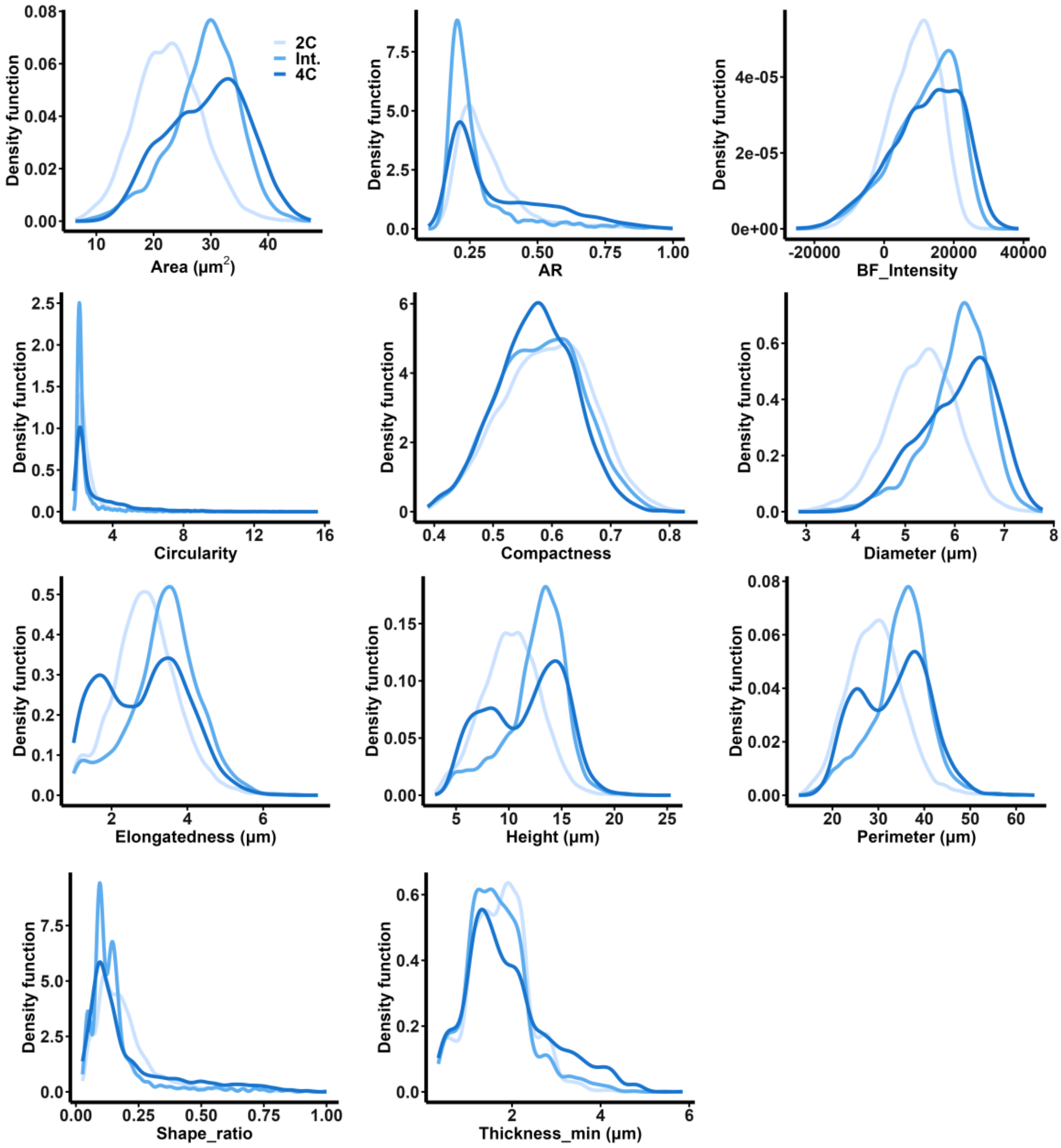

2C
Int.
4C
Density function
Area (μm2)
AR
BF_Intensity
Circularity
Compactness
Diameter (μm)
Elongatedness (μm)
Height (μm)
Perimeter (μm)
Shape_ratio
Thickness_min (μm)


### S5 Fig

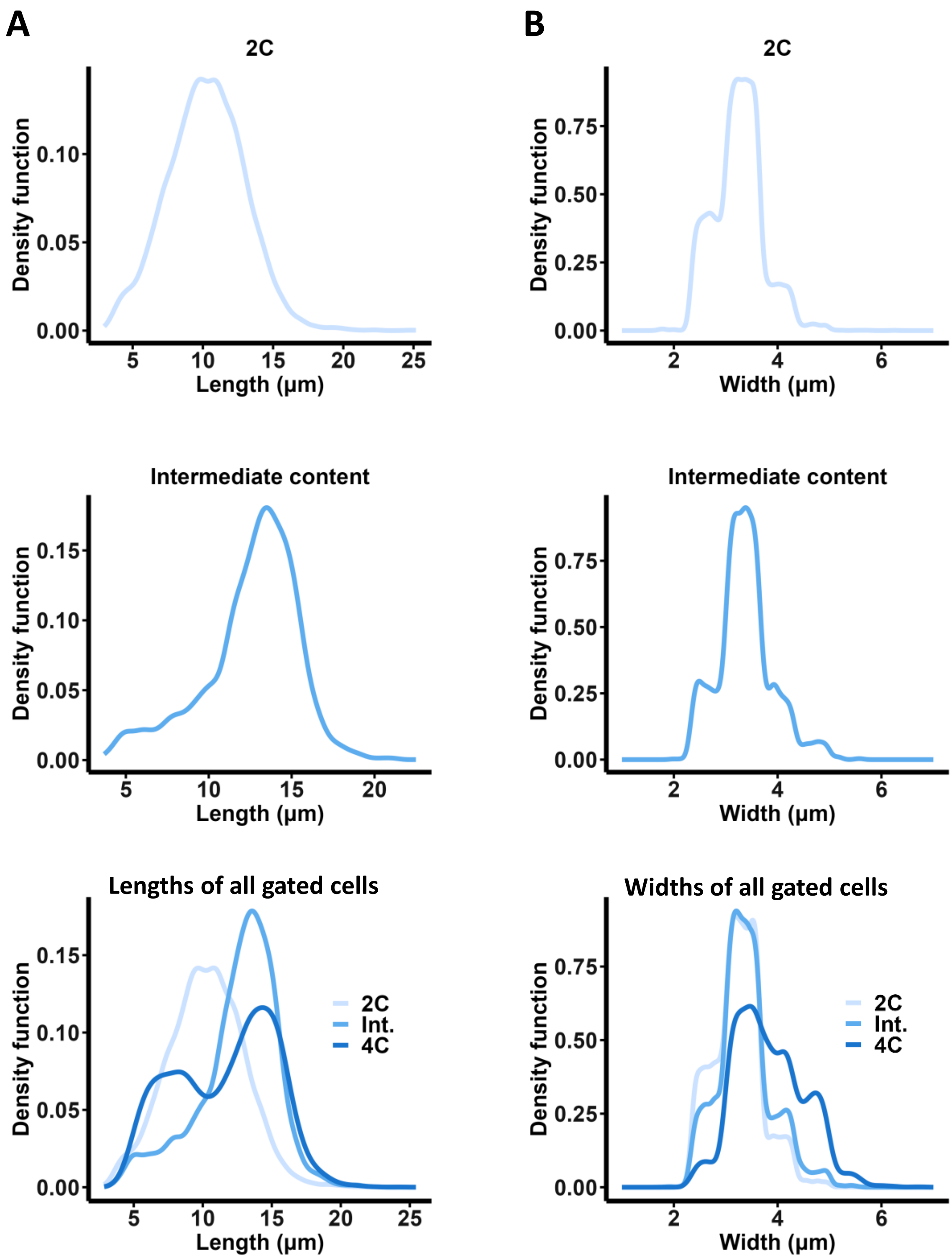

A
2C
Density function
0.10
0.05
0.00
5
10
15
20
25
Length (µm)
B
2C
Density function
0.75
0.50
0.25
0.00
2
4
6
Width (µm)
Intermediate content
Density function
0.15
0.10
0.05
0.00
5
10
15
20
Length (µm)
Intermediate content
Density function
0.75
0.50
0.25
0.00
2
4
6
Width (µm)
Lengths of all gated cells
Density function
0.15
0.10
0.05
0.00
2C
Int.
4C
5
10
15
20
25
Length (µm)
Widths of all gated cells
Density function
0.75
0.50
0.25
0.00
2C
Int.
4C
2
4
6
Width (µm)


### S6 Fig

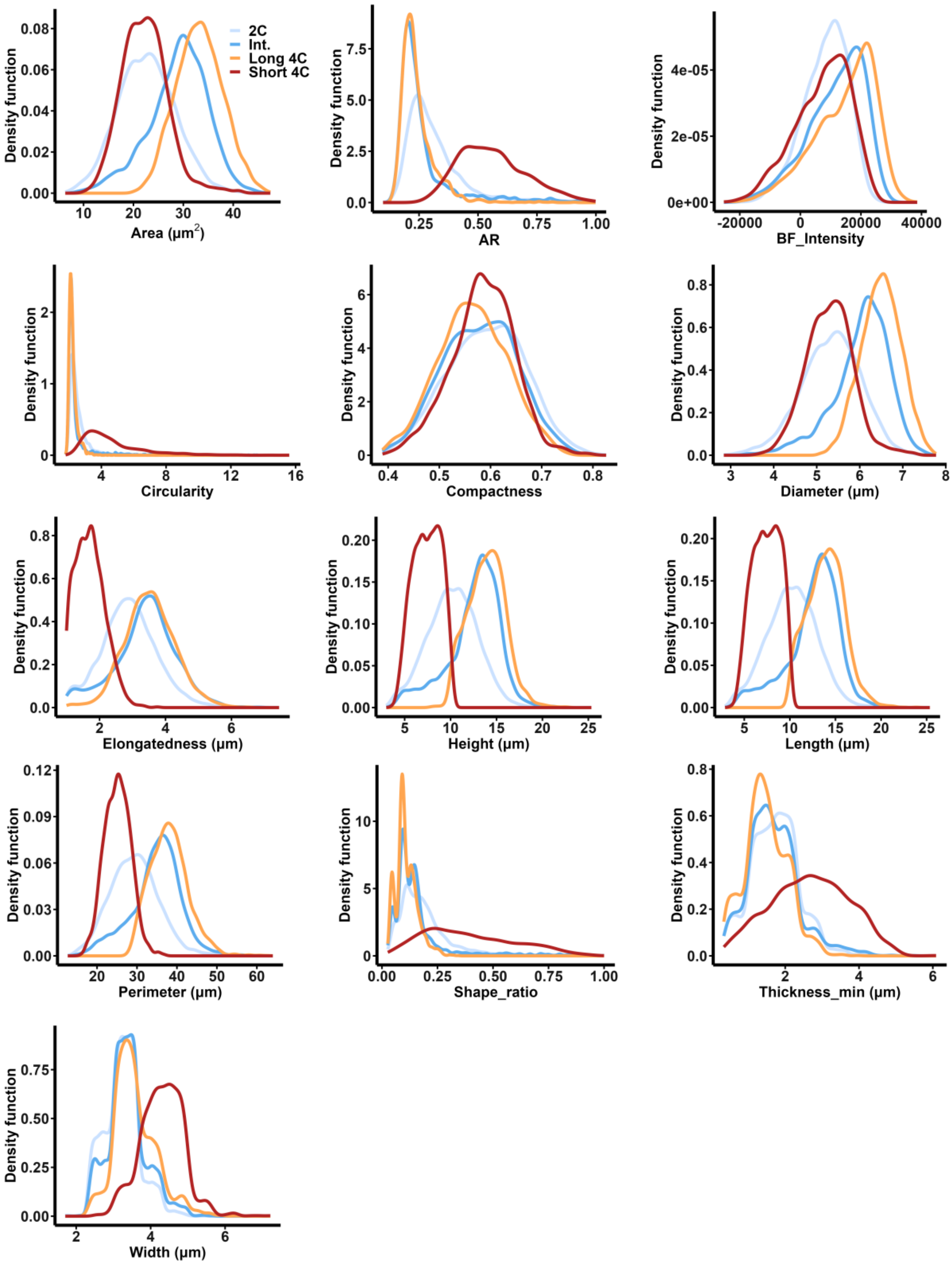

2C
Int.
Long 4C
Short 4C
Density function
Area (μm2)
AR
BF_Intensity
Circularity
Compactness
Diameter (μm)
Elongatedness (μm)
Height (μm)
Length (μm)
Perimeter (μm)
Shape_ratio
Thickness_min (μm)
Width (μm)


**S7 Fig**

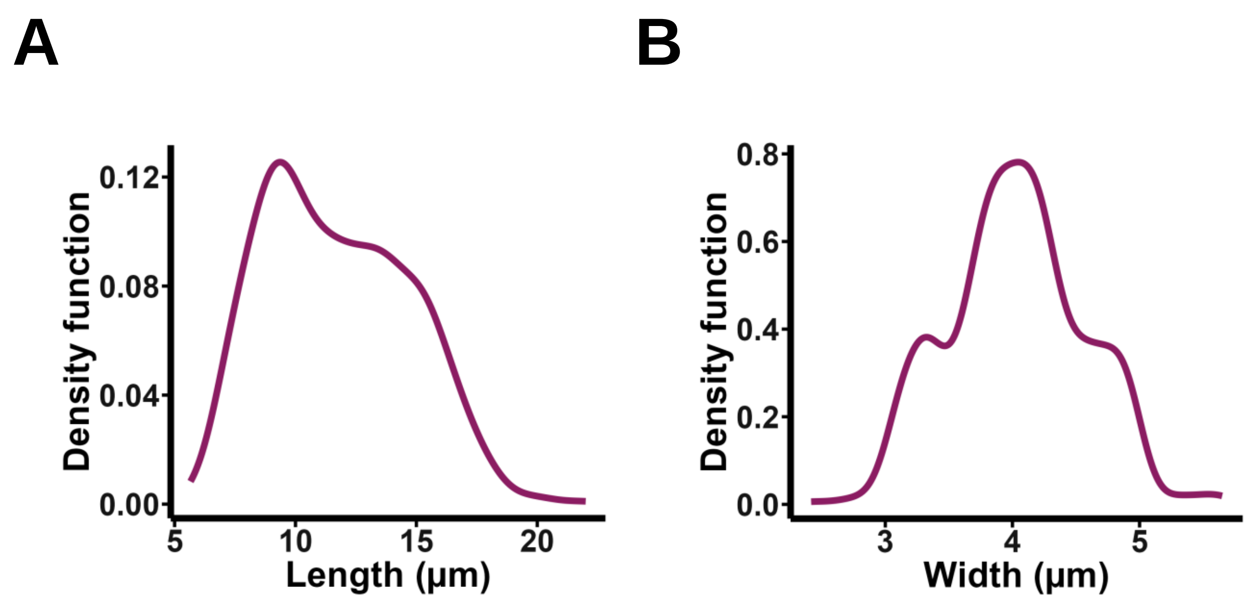
A
Density function
0.12
0.08
0.04
0.00
5
10
15
20
Length (μm)
B
Density function
0.8
0.6
0.4
0.2
0.0
3
4
5
Width (μm)

S8 Fig

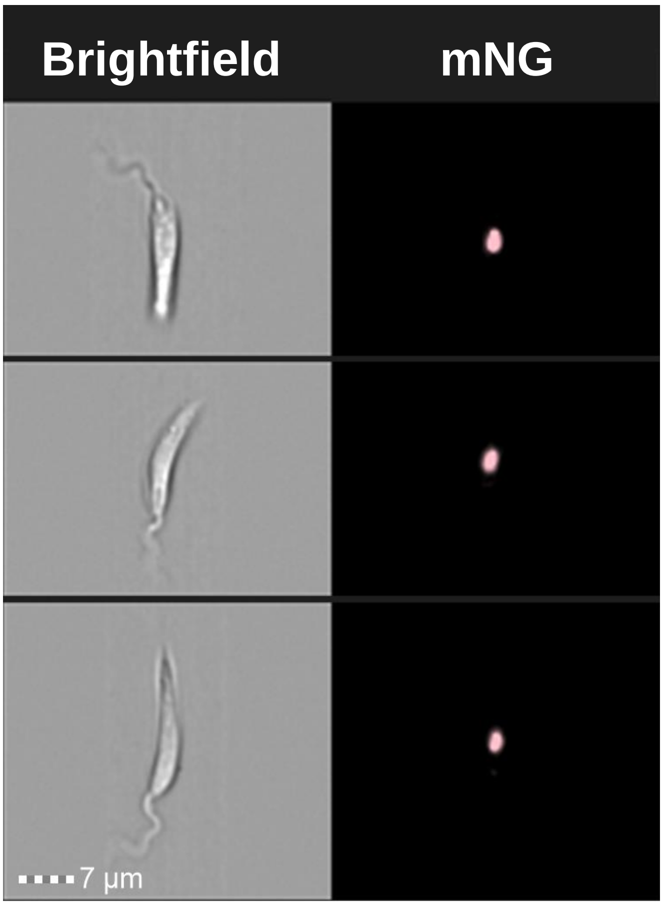


**S9 Fig**

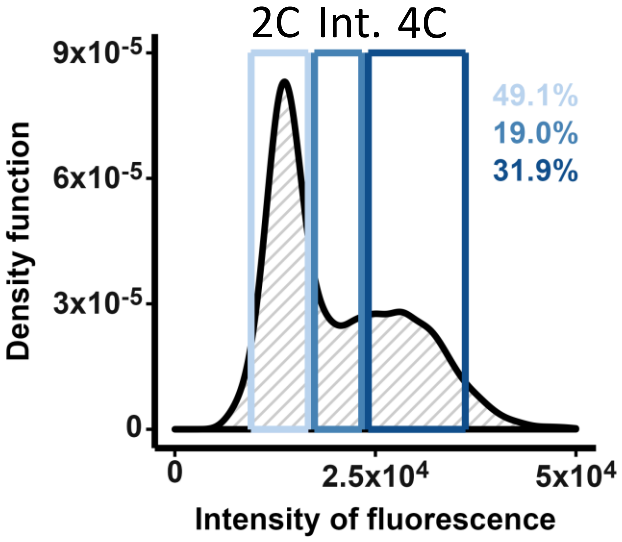

2C
Int.
4C
9x10-5
6x10-5
3x10-5
0
Density function
49.1%
19.0%
31.9%
0
2.5x10^4
5x10^4
Intensity of fluorescence


**S10 Fig**

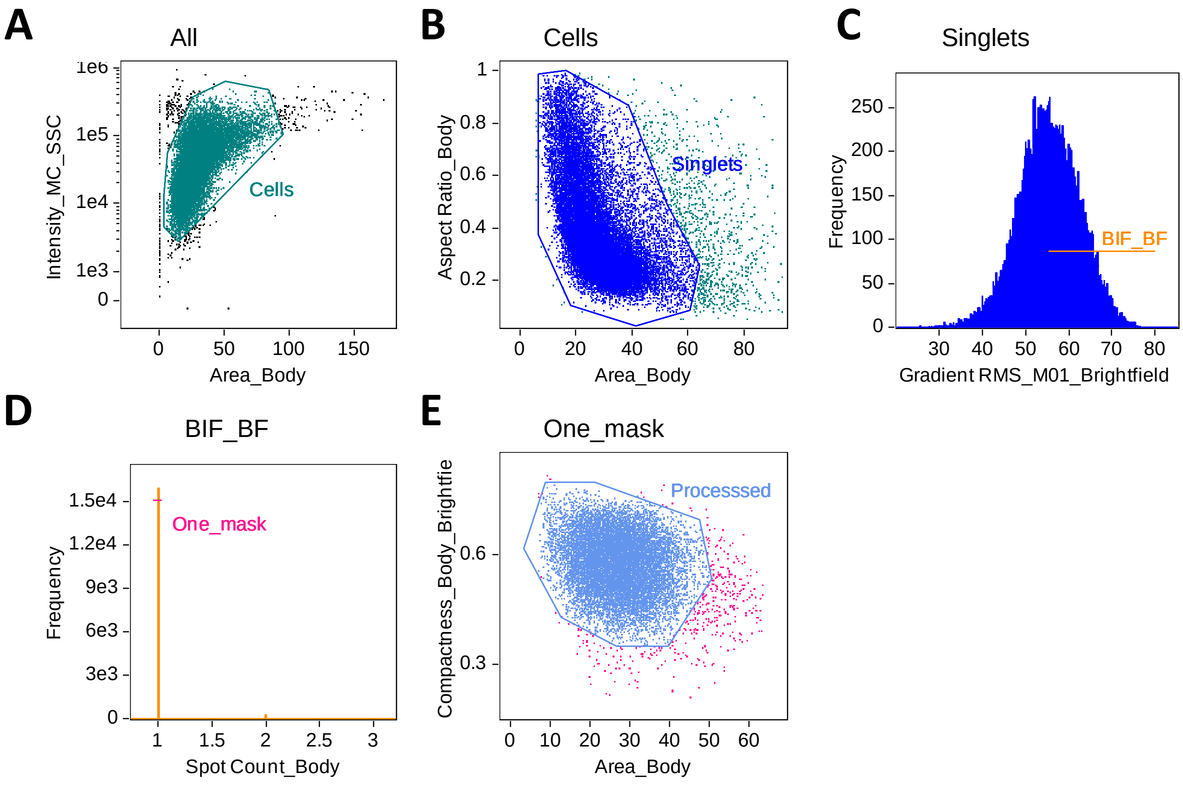
A
All
Intensity_MC_SSC
Area_Body
Cells
B
Cells
Aspect Ratio_Body
Area_Body
Singlets
C
Singlets
Frequency
Gradient RMS_M01_Brightfield
BIF_BF
D
BIF_BF
Frequency
Spot Count_Body
One_mask
E
One_mask
Compactness_Body_Brightfie
Area_Body
Processsed

### S11 Fig

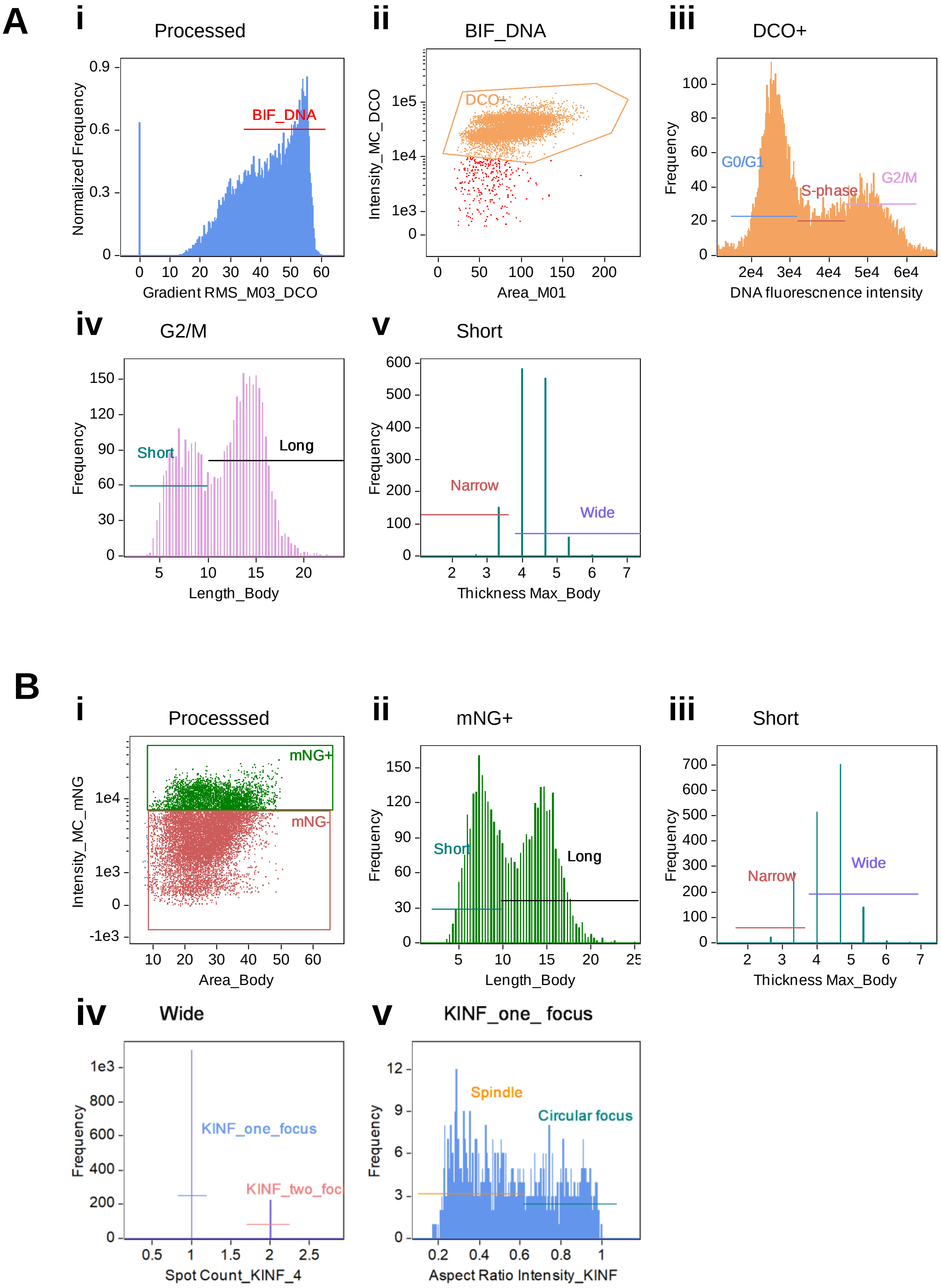
A
i
Processed
Normalized Frequency
BIF_DNA
Gradient RMS_M03_DCO
ii
BIF_DNA
Intensity_MC_DCO
DCO+
Area_M01
iii
DCO+
Frequency
G0/G1
S-phase
G2/M
DNA fluorescnence intensity
iv
G2/M
Frequency
Short
Long
Length_Body
v
Short
Frequency
Narrow
Wide
Thickness Max_Body
B
i
Processsed
Intensity_MC_mNG
mNG+
mNG-
Area_Body
ii
mNG+
Frequency
Short
Long
Length_Body
iii
Short
Frequency
Narrow
Wide
Thickness Max_Body
iv
Wide
Frequency
KINF_one_focus
KINF_two_foc
Spot Count_KINF_4
v
KINF_one_ focus
Frequency
Spindle
Circular focus
Aspect Ratio Intensity_KINF

### S12 Fig

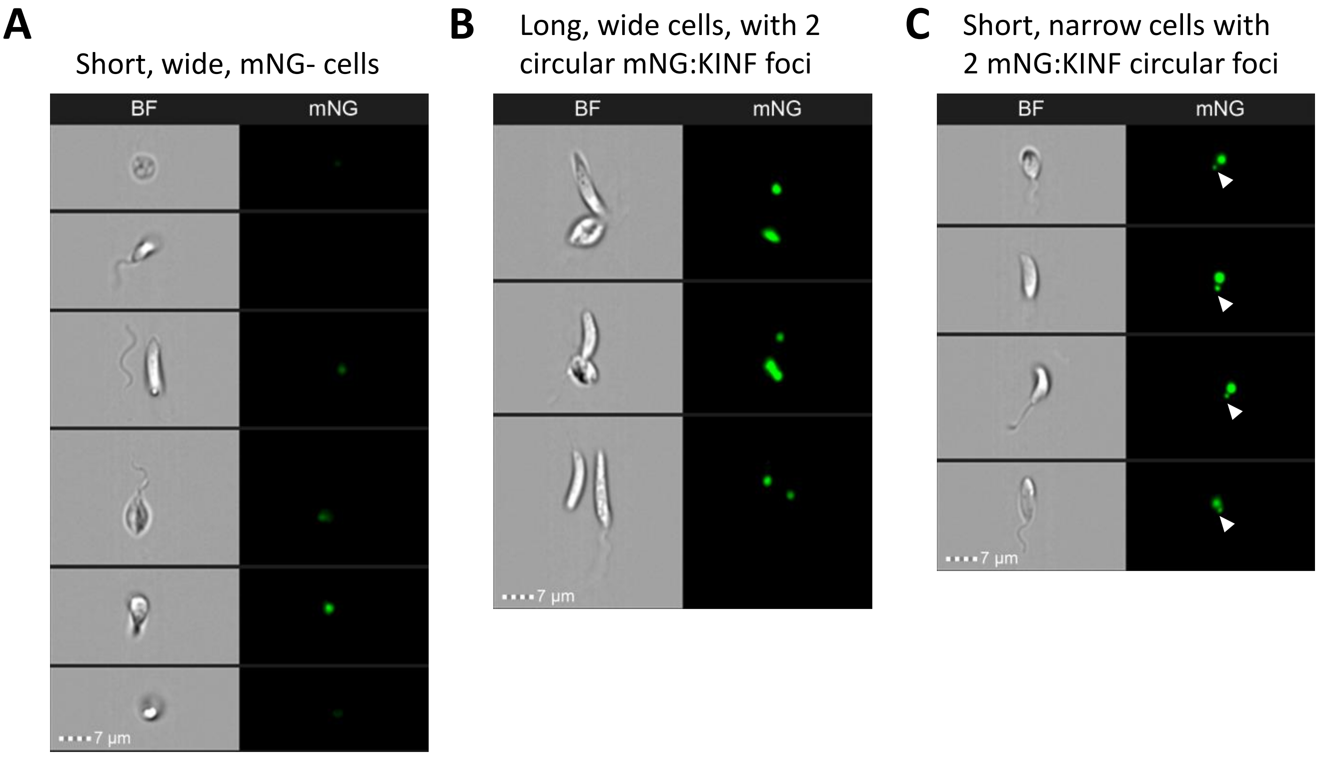
A
Short, wide, mNG- cells
BF
mNG
7 µm
B
Long, wide cells, with 2 circular mNG:KINF foci
BF
mNG
7 µm
C
Short, narrow cells with 2 mNG:KINF circular foci
BF
mNG
7 µm

### S13 Fig
